# Non-invasive auditory brainstem responses to FM sweeps in awake big brown bats

**DOI:** 10.1101/2022.06.03.494657

**Authors:** Andrea Megela Simmons, Amaro Tuninetti, Brandon M. Yeoh, James A. Simmons

## Abstract

We introduce two EEG techniques, one based on conventional monopolar electrodes and one based on a novel tripolar electrode, to record for the first time auditory brainstem responses (ABRs) from the scalp of unanesthetized, unrestrained big brown bats. Stimuli were frequency-modulated (FM) sweeps varying in sweep direction, sweep duration, and harmonic structure. As expected from previous invasive ABR recordings, upward-sweeping FM signals evoked larger amplitude responses (peak-to-trough amplitude in the latency range of 3-5 ms post-stimulus onset) than downward-sweeping FM signals. Scalp-recorded responses displayed amplitude-latency trading effects as expected from invasive recordings. These two findings validate the reliability of our noninvasive recording techniques. The feasibility of recording noninvasively in unanesthetized, unrestrained bats will energize future research uncovering electrophysological signatures of perceptual and cognitive processing of biosonar signals in these animals, and allows for better comparison with ABR data from echolocating cetaceans, where invasive experiments are heavily restricted. Because experiments can be repeated in the same animal over time without confounds of stress or anesthesia, our technique requires fewer captures of wild bats, thus helping to preserve natural populations and addressing the goal of reducing animal numbers used for research purposes.

## Introduction

Echolocating bats exhibit extremely fine temporal acuity in perception as an integral part of auditory signal processing for echoes they receive from their environments (Simmons et al. 2014). The neural mechanisms underlying this remarkable perceptual feat are topics of great scientific and technological interest (Ballieri et al. 2017). A variety of invasive electrophysiological techniques have been employed to decipher the neural bases of echolocation in bats, including auditory brainstem responses (ABRs), local field potentials, intracellular and extracellular recordings from single or multiple neurons, and iontophoresis of neuropharmacological agents (reviews: Pollak and Casseday 2012; Faure and Firzlaff 2016; Pollak 2016). Newer methods, including multiple-electrode arrays for recording extracellular activity in stationary (Luo et al. 2019) and free-flying bats (Kothari et al. 2018), and calcium imaging combined with extracellular multiple neuron recordings (Simmons et al. 2020), have been introduced. All of these techniques provide essential information on brain representation of biosonar signals and echoes, but all are constrained by requiring anesthesia for invasive surgical preparation and, in most experiments, during neural recordings as well. Even experiments using subcutaneous needle electrodes to record ABRs require bats to be anesthesized during both needle insertion and data collection (Burkard and Moss 1994; Linnenschmidt and Wiegrebe 2019; Lattenkamp et al. 2021; Möckel et al. 2021). The use of anesthesia precludes direct analysis of the bat’s percepts or cognitive processing of acoustic signals. Moreover, US federal regulations require animals to be euthanized after reaching experimental endpoints, which are often defined as at the termination of individual electrophysiological sessions. The ability to record electrophysiological activity noninvasively from awake behaving bats would advance the goals of unraveling the mechanisms of the bat’s fine temporal acuity, while introducing a refinement of experimental technique leading to reduction in numbers of experimental animals required for experiments. Such reductions will help maintain local endangered bat populations.

Echolocation in bats shares fundamental similarities with echolocation in dolphins and may be based on similar signal processing algorithms (Au and Simmons 2007; Branstetter et al. 2007; Ming et al. 2021). The big brown bat (*Eptesicus fuscus*) and the bottlenose dolphin (*Tursiops truncatus*) demonstrate similar microsecond-scale acuity to echo jitter and phase (Simmons 1979, 1993; Simmons et al. 1990; Finneran et al. 2019, 2020), and to fine spectral features of echoes (Shriram and Simmons 2019; Accomando et al. 2020). The search for shared neural mechanisms underlying this sensitivity has been limited by the availability of correspondingly shared techniques. Because dolphins, like all cetaceans, are protected species, invasive recordings for research purposes are highly restricted and thus not commonly employed (but see Bullock et al. 1968). Instead, our understanding of brain function in these animals relies heavily on data from non-invasive functional imaging and electrophysiological (including ABR; auditory evoked potentials; electroencephalography, EEG) recordings from unanesthetized animals using surface or subcutaneous electrodes (Ridgway et al. 1981, 2006; Nachtigall and Schuller 2014; Schalles et al. 2021). Because of these methodological differences, particularly related to the effects of anesthesia on brain function, direct comparisons of electrophysiological data from bats and from dolphins can be challenging.

In this paper, we introduce non-invasive EEG techniques for recording brain activity from awake, unrestrained big brown bats. These techniques require training of bats to remain motionless, thus eliminating the need for restraint, but neither anesthesia nor surgery. Because our methods are completely noninvasive, the same bats can continue to participate in experiments over days or weeks without any risk from stress, anesthesia, surgical or electrode damage to the brain, or systemic infection. In the absence of these complicating factors, the same animals can still participate in behavioral studies as approved in research protocols.

Previously, we (Luo et al. 2019) demonstrated that short-latency ABRs recorded invasively from the big brown bat’s inferior colliculus showed a similar response pattern to frequency-modulated upsweeps (FM-up) and downsweeps (FM-down) as those recorded non-invasively from the bottlenose dolphin’s scalp (Finneran et al. 2017): Responses to FM-upsweeps are larger in amplitude compared to responses to FM-downsweeps. Similar patterns of response in both dolphins and humans have been interpreted as providing estimates of traveling wave velocities and insight into basilar membrane operation (Dau et al. 2000; Elberling et al. 2007; Finneran et al. 2017). Here, we employ these same stimuli to compare the effects of sweep direction on scalp recordings with those observed in invasive recordings, and we use these comparisons to evaluate our new noninvasive technique. Replication of previous results with a non-invasive method will energize additional studies validating results from other invasive experiments.

## Methods

### Animals

Two adult female big brown bats (J and T, ages unknown) participated in these experiments. They were captured from local barns, as authorized by a State of RI scientific collection permit. To conserve local bat populations, the state permit limits severely the numbers of animals that can be captured in any given year and thus restricts numbers on-hand in the laboratory for research purposes. Bats were socially-housed in the laboratory in a wire frame enclosure (6’ x 8’ x 8’) within a larger colony room. They were vaccinated for rabies and individually identified by readable microchips inserted under the skin of their backs (Trovan ID-100A RFID transponder, Trovan LID-573 microchip reader). They had unlimited access to fresh water and were provided daily with sufficient vitamin-enriched live mealworms *(Tenebrio larvae)* to keep their body weights within a healthy range of 15-20 g. Bats were allowed to fly unfettered in a large flight room for weekly exercise; both of them echolocated and flew without difficulty. Both bats also emitted communication and echolocation sounds in their home enclosure (monitored by Wildlife Acoustics recording devices). The colony room was maintained at temperatures of 20-24° C and 55-65% relative humidity. All experimental and husbandry procedures were approved by the Brown University Institutional Animal Care and Use Committee and adhere to US federal guidelines.

As the first step in our procedure, the two bats were trained to sit without excessive movements for periods up to 30 min in a 50-mm deep ceramic dish, with their torsos within the dish and their heads resting on the edge. The dish was placed on an elevated steel platform in a single-walled sound-attenuating, electrically-shielded recording booth (Industrial Acoustics Co, N. Aurora IL). Training took place over several weeks. At the beginning of training, a bat was placed inside the ceramic dish; this ceramic dish is familiar to the bats, being the same as the food dishes used in their social housing. In the colony room, bats often rest or sleep in their food dishes after eating their daily food allotment. If the bat began crawling out of the dish on the platform, a soft broadband “shh” sound was made by the trainer to indicate to the bat that it had made an error, then the bat was gently placed back into the dish. If the bat remained inside the bowl for 20-30 consecutive sec, it was rewarded with a mealworm, which it ate inside the dish. This was repeated multiple times, with rewards given every 30 sec, until the bat stayed in the dish for a period of 2 consecutive min. At this time point, a single electrode covered in conductive paste was placed on the bat’s back. If the bat tolerated the electrode placement without excessive movement, it was given another mealworm reward immediately after. If the bats reacted by leaving the dish, a “shh” error signal was again made by the trainer, and the electrode was reattached once the bat was back in the dish and had settled down. If the bat tolerated attachment of the first electrode, a second electrode was placed on its back, and then the third electrode placed along the midline of its scalp. A mealworm reward was provided after application of each of the first two electrodes, but not the third, as we wished to avoid excessive head movement. Once all three electrodes were applied to the bat, ultrasonic FM stimuli were emitted towards the bat at full amplitude and at the same repetition rate as used in data collection. Four individual bats began training in this manner, but two were removed after showing no behavioral progress after several days of training. Training was conducted 2-3 times a week for 2-3 weeks, until the two remaining bats could reliably tolerate having three electrodes applied with conductive paste while sitting motionless in the dish, while exposed to repeating auditory stimuli at full amplitude, for at least 5 consecutive min. The two bats used in this experiment learned to sit motionless in the dish for up to 30 min. Once the bats reached the 5 min criterion, the hair on their heads and their lower back was trimmed down with safety scissors, and diluted depilatory (Nair™; Church & Dwight, Ewing NJ) was applied for 2 min to remove remaining hair. Because recording electrodes will not adhere to hair and because bats’ hair grew back within a few days after depilation, hair removal was repeated as necessary during the time frame of these experiments. Bats tolerated the hair removal procedure well, without any signs of skin irritation or permanent loss of hair.

### Acoustic stimulation

Acoustic stimuli were generated as digital .wav files using Adobe Audition v. 12.1 (Adobe Inc, San Jose CA) at a sampling rate of 500 kHz. They consisted of FM-up and FM-downsweeps, containing one or two harmonics, at durations of 3, 2, 1, 0.7, 0.5, 0.3, 0.2, and 0.1 ms (**Fig. 1**). FM sweeps with a single harmonic (FM-1H) covered the frequency range of 20-100 kHz; FM sweeps with two harmonics (FM-2H) covered the frequency range of 20-50 kHz in the first harmonic and 40-100 kHz in the second harmonic. All stimuli had raised cosine envelopes with 50% rising and falling shapes, matching the envelope shape of signals used by Luo et al. (2019). The natural FM echolocation sounds of big brown bats are two harmonic FM-downsweeps varying in duration from about 15 ms to about 0.6 ms over a pursuit/capture sequence, depending on the surrounding environment (Surlykke and Moss 2000). Stimulus durations chosen for this experiment fall within the shorter ends of this biological range and again mimicked those used for invasive recordings (Luo et al. 2019).

**Fig 1.**
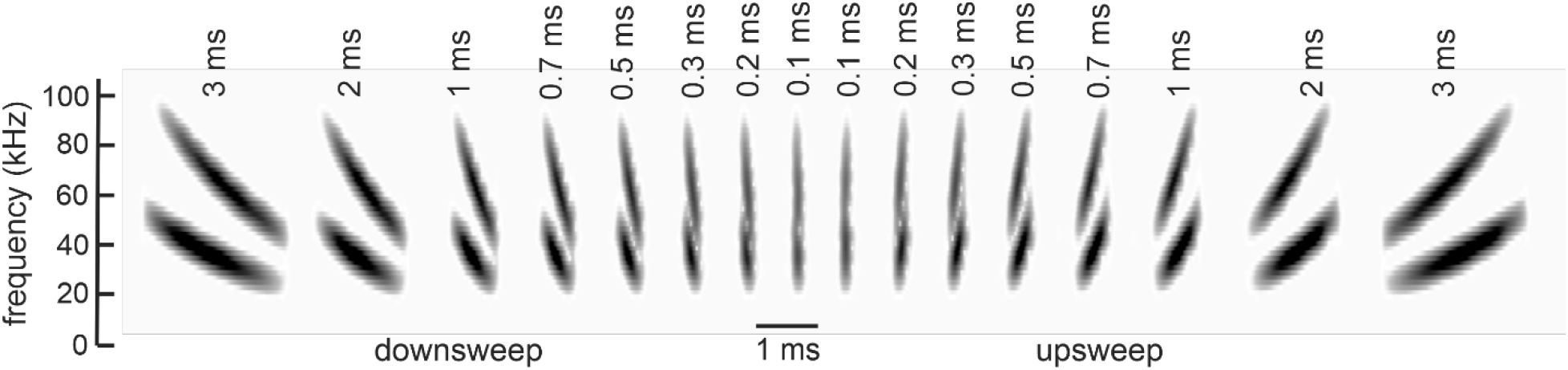
Spectrograms of FM-2H logarithmic down- and up-sweeping FM sweeps at the bat’s midline, after passing through the TDT tweeter. Sweeps varied in duration. FM-1H stimuli (not shown) did not include the second harmonic.

The rationale for presenting short FM sweeps varying in sweep direction and duration derives from models of basilar membrane mechanics in other mammals (Robles and Ruggero 2001). FM-downsweeps, such as those used by the bat for echolocation, contain high frequencies followed by low frequencies. The traveling wave’s direction of propagation from high frequencies at the base of the cochlea to low frequencies at the apex causes low frequencies to be delivered to their receptors slightly later than those of high frequencies. The consequence is a delayed activation of eighth nerve fibers tuned to low frequencies compared to those tuned to high frequencies. FM-upsweeps contain low frequencies followed by high frequencies. In this case, low frequencies travel to their maximal place of excitation towards the apex before high frequencies arrive at their maximal place of excitation towards the base. Here, the delayed activation of low frequency responses is counteracted. FM sweeps thus produce an asymmetric pattern of excitation, visualized as the time-place appearance of different frequencies according to the direction of the sweep (up *versus* down) and whether the sweep adds to the time-delay of frequencies at successive places (for downsweeps) or counteracts the time-delays (for upsweeps). ABRs reflect synchronous neural activity in the ascending auditory pathway. When eighth nerve responses are dispersed in time, as in an FM-downsweep, response synchrony is weaker, leading to a lower amplitude ABR, than if all frequencies arrived at their tuned location simultaneously. At appropriate durations of an FM-upsweep, neural responses at all frequencies can be brought into alignment and summate to produce a stronger synchronous response (Dau et al. 2000). The optimal duration of an FM-upsweep that counteracts frequency dispersion along the basilar membrane and produces this stronger response is predicted to match the velocity of the travelling wave (Elberling et al. 2007). In bottlenose dolphins, the optimal FM-upsweep duration derived from surface ABR recordings lies within the range of 0.45-1.1 ms, depending on stimulus level (Finneran et al. 2017); in big brown bats, the optimal FM-upsweep duration derived from invasive recordings lies within the range of 0.5-1 ms (Luo et al. 2019).

Digitized acoustic stimuli were stored on a Dell Windows 10 laptop computer located outside the recording booth for call-up during an experiment. Stimuli were presented through an ultrasonic tweeter loudspeaker placed 45 cm away from and facing towards the midline of the bat’s head. In initial experiments, we presented sounds through a Kenwood high-frequency KFC-XT15ie tweeter loudspeaker (Kenwood Corp, Tokyo JP). The frequency response of the Kenwood tweeter varied +2 to −9 dB across the frequency range of 20 to 90 kHz and decreased by 25 dB at 100 kHz. In later experiments, we presented sounds through a TDT electrostatic speaker (ES1 speaker driven by an ED1 speaker driver; Tucker-Davis Technologies, Alachua FL), because of its better higher frequency response (±9 dB over the frequency range 4-110 kHz (Table 1). As used here, the maximum output of the Kenwood tweeter was 98 dB peSPL at 20-60 kHz, and that of the TDT tweeter was 90 dB peSPL at 25-80 kHz. Stimulus levels for both tweeters were calibrated by placing a Brüel & Kjaer Model 4135 (“¼”) condenser microphone at the position occupied by the bat’s head during experiments. Stimulus amplitude is expressed as dB peSPL re 20 μPa. During experiments, the acoustic stimuli delivered to the bat were monitored using a Dodotronic Momimic ultrasonic microphone (Dodotronic, Castel Gandolfo IT) suspended over the bat’s head, whose output was connected to one channel of a Tektronix Type 2000 70 MHz 4-channel digital oscilloscope (Tektronix Inc, Beaverton OR) located outside the recording booth. Experimental parameters on each recording day are listed in Table 1.

**Table 1.**
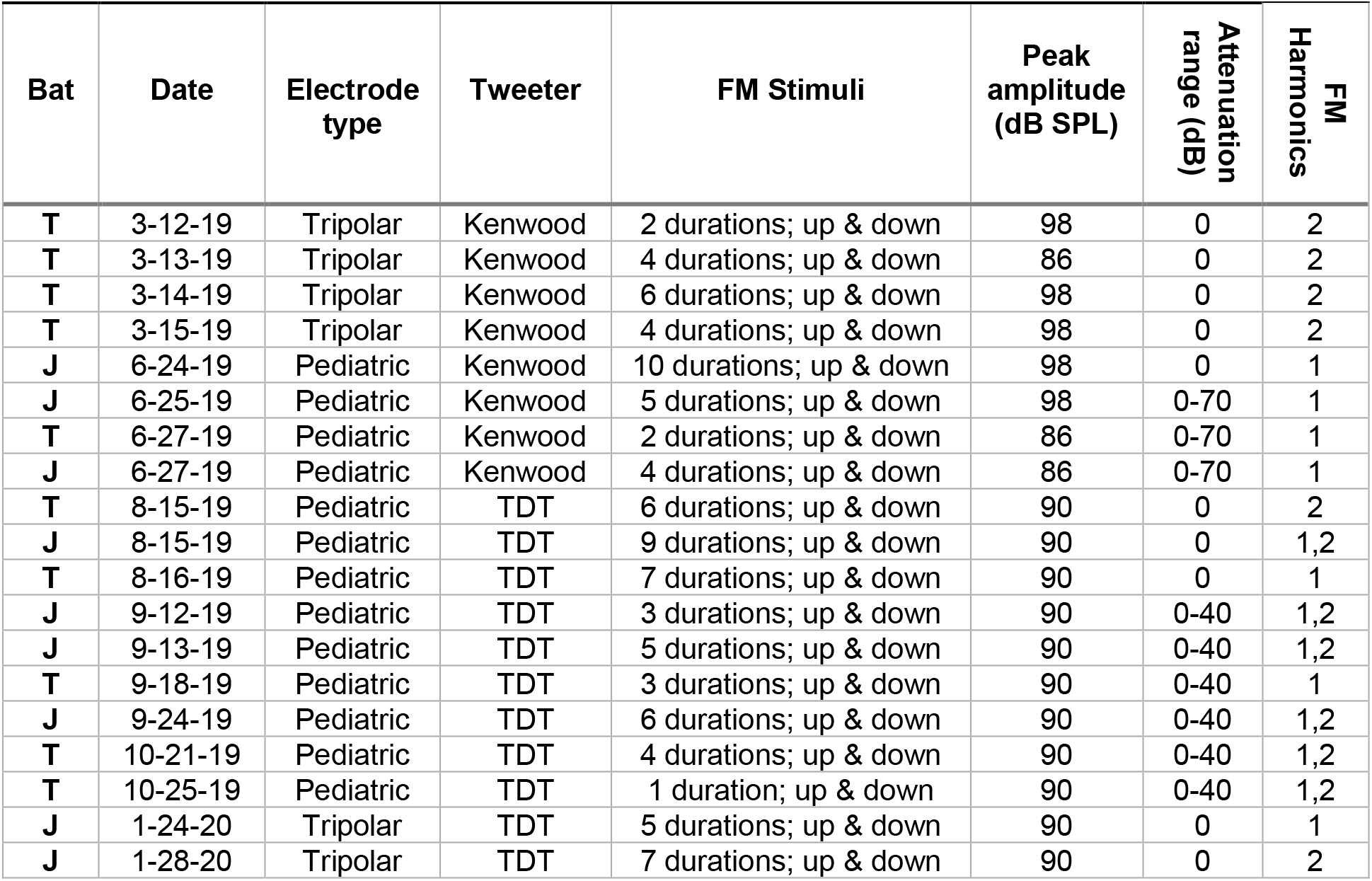
Experimental parameters.

| Bat | Date | Electrode type | Tweeter | FM Stimuli | Peak amplitude (dB SPL) | Attenuation range (dB) | FM Harmonics |
| --- | --- | --- | --- | --- | --- | --- | --- |
| T | 3-12-19 | Tripolar | Kenwood | 2 durations; up & down | 98 | 0 | 2 |
| T | 3-13-19 | Tripolar | Kenwood | 4 durations; up & down | 86 | 0 | 2 |
| T | 3-14-19 | Tripolar | Kenwood | 6 durations; up & down | 98 | 0 | 2 |
| T | 3-15-19 | Tripolar | Kenwood | 4 durations; up & down | 98 | 0 | 2 |
| J | 6-24-19 | Pediatric | Kenwood | 10 durations; up & down | 98 | 0 | 1 |
| J | 6-25-19 | Pediatric | Kenwood | 5 durations; up & down | 98 | 0-70 | 1 |
| T | 6-27-19 | Pediatric | Kenwood | 2 durations; up & down | 86 | 0-70 | 1 |
| J | 6-27-19 | Pediatric | Kenwood | 4 durations; up & down | 86 | 0-70 | 1 |
| T | 8-15-19 | Pediatric | TDT | 6 durations; up & down | 90 | 0 | 2 |
| J | 8-15-19 | Pediatric | TDT | 9 durations; up & down | 90 | 0 | 1,2 |
| T | 8-16-19 | Pediatric | TDT | 7 durations; up & down | 90 | 0 | 1 |
| J | 9-12-19 | Pediatric | TDT | 3 durations; up & down | 90 | 0-40 | 1,2 |
| J | 9-13-19 | Pediatric | TDT | 5 durations; up & down | 90 | 0-40 | 1,2 |
| T | 9-18-19 | Pediatric | TDT | 3 durations; up & down | 90 | 0-40 | 1 |
| J | 9-24-19 | Pediatric | TDT | 6 durations; up & down | 90 | 0-40 | 1,2 |
| T | 10-21-19 | Pediatric | TDT | 4 durations; up & down | 90 | 0-40 | 1,2 |
| T | 10-25-19 | Pediatric | TDT | 1 duration; up & down | 90 | 0-40 | 1,2 |
| J | 1-24-20 | Tripolar | TDT | 5 durations; up & down | 90 | 0 | 1 |
| J | 1-28-20 | Tripolar | TDT | 7 durations; up & down | 90 | 0 | 2 |

All electronic equipment needed for sound presentation and monitoring was located outside the recording booth, connected by cables running through small openings in the booth wall. For playback during experiments, the stored acoustic stimuli were uploaded from the Dell computer to a Koolertron 15-mHz DDS Signal Generator device (500 kHz digital-to-analog sampling rate; Model GH-CJDS66-A, Shenzhen Kuleton Technology, Shenzhen China). The DDS Signal Generator device output was triggered by a Biopac MP160 data acquisition system with AcqKnowledge 5 software (Biopac Systems, Goleta CA). The analog output of the DDS device was routed into a TDT PA5 attenuator (Tucker-Davis Technologies, Alachua FL), a Harman-Kardon PM545 stereo power amplifier (Harman International Industries, Stamford CT), and finally to the tweeter inside the recording booth. The electronic trigger for producing each sound, the electrical waveform delivered to the power amplifier, and the physiological signal recorded from the bat (see below) were recorded on the other three channels of the 4-channel Textronix oscilloscope located outside the recording booth.

### Electrodes and recording set-up

Two different types of electrodes were used for recording brain activity from the bat’s scalp. One type was a Natus silver-cup 6-mm diameter conventional monopolar pediatric EEG electrode (pEEG; SKU 019-772100, MVAP Medical Supplies, Thousand Oaks CA). An electrode was placed on the posterior scalp using conductive electrode paste (Weaver Ten20™; Weaver & Co, San Diego CA). A second and third electrode were placed posterior, on the bat’s bare upper and lower back, for differential recording and grounding. The second type was a novel tripolar EEG electrode (Besio et al. 2006; CREmedical, Kingston RI), consisting of three concentric conductive contact rings—an outer ring with a diameter of 6 mm, an intermediate-sized contact ring, and a central contact point. Tripolar EEG recordings (tEEG) are based on a nested sequence of differential stages that extract voltage differences between the outer and intermediate rings, the intermediate ring and central contact, and the two rings to the central contact (Besio et al. 2006). The final output is the voltage attributable to the central contact alone, which results in suppression of common-mode artifacts not only from muscles but also from regions of the brain remote from the central contact. The tripolar electrode produces one differential signal via a custom preamplifier (CREmedical, Kingston RI), with its differential analog tripolar output joined to the overall ground electrode for a second differential stage with a gain of 20X. This CRE preamplifier was located on the floor inside the recording booth, and then remotely connected to the rest of the electrophysiological equipment outside the booth. The analog outputs of the pEEG and the tEEG signals were remotely connected to a Biopac MP160 System and ERS100C evoked-response hardware module (Biopac Systems, Goleta CA) for averaging, analog-to-digital conversion, and subsequent digital signal-processing. The ERS100C module was set to filters from 100-20,000 Hz and included a gain of 50,000X.

Electrophysiological responses were acquired in repetitive segments of 50 ms (the acquisition window length) triggered by stimulus presentations, digitized at a sampling frequency of 10 kHz and subsequently added to the ongoing averaged electrophysiological response. A real-time display of the building-up and averaging of the response was programmed into the Biopac display on its host computer (Dell Windows 10 Laptop connected via USB for operation with the AcqKnowledge 5 program). Recorded signals were then saved as .txt files.

### Procedure

Bats were allowed to position themselves inside the ceramic dish until they rested comfortably. Both bats rested their heads on the rim of the dish, which was rotated if necessary to point directly towards the loudspeaker position. Either pEEG or tEEG electrodes were applied to the bat’s exposed posterior scalp with conductive paste. We recorded bats’ body temperatures before and after each recording session; temperature never varied by more than 4° C before these two time points. If bats moved excessively during recordings, we offered them sips of water or pieces of mealworms and short breaks (during which electrodes were often reapplied). Recording sessions lasted 5-30 min, depending on the bat’s ability to remain relatively motionless and the quality of the evoked response. Differences in session length resulted in uneven sample sizes for the different stimulus types.

A FM-1H 1 ms duration upsweep at 98 dB peSPL (Kenwood tweeter) or 90 dB peSPL (TDT tweeter) was presented as a search stimulus to assess whether electrode placements yielded stable baseline and high-amplitude evoked activity. Once a good electrode site was identified, we then presented stimuli varying in harmonic structure (1H or 2H), sweep duration, and sweep direction (Table 1), with the order determined by a random number generator, at a rate of 3.2/s for 200 repetitions (on two recording days with high background noise levels, repetitions were increased to 500). Stimulus levels were set at 86 or 98 dB peSPL (Kenwood tweeter) or at 90 dB peSPL (TDT tweeter). An interval of 8-30 s (average of 10 s; long enough to save data files and upload a new sound file) separated presentations of different stimuli. In other experiments (Table 1), we presented FM sweeps at a range of levels, decreasing in steps of 10 dB. This manipulation allowed us to evaluate the presence of amplitude-latency trading, a feature of neural responses prevalent in invasive recordings (Pollak 1988; Simmons et al. 1990; Klug et al. 2000). We hypothesized that amplitude-latency trading would be observed in surface ABRs as well. At the end of each recording session, bats were given water and mealworms, and returned to their home cages.

### Data processing

Electrophysiological responses were averaged over the 200 or 500 stimulus presentations while being visualized in realtime, and then saved to disk unless the recording was disrupted by bat movement. These files were imported as .txt files into MATLAB 2019b (MathWorks, Natick MA) for processing using custom scripts. Each response was demeaned to remove DC offset and filtered by a 48th order linear phase bandpass FIR filter with cutoff frequencies of 300-3000 Hz. A threshold of three times the RMS value of activity in the first 1 ms of the response, consistent with the threshold used by Luo et al. (2019), was set to identify evoked responses from baseline activity. Positive and negative peaks in the averaged response were identified automatically using the *findpeaks* function in MATLAB, and then confirmed visually. We quantified the amplitude and latency of the largest positive peak and the subsequent negative peak within a specific, short latency range (see below) to calculate peak-to-trough amplitudes. In experiments where stimulus level was varied, we first quantified these metrics at the highest stimulus level presented, then traced any changes in amplitude or latency of this peak-to-trough response at progressively lower sound levels. Response latency was calculated from stimulus onset at the bat’s ears, taking into consideration the acoustic delay of 1.25 ms produced by the 45 cm distance between the bat and the tweeter. Responses were also visualized as heat maps in which warmer colors indicate higher positive amplitudes and bluer colors indicate lower amplitudes.

## Results

Data were collected from 13 pEEG recording sessions and 6 tEEG recording sessions from the two bats (Table 1). Because the bat’s ability to remain motionless was the main experimental constraint, recording sessions varied in length and the entire stimulus set could not be presented in all sessions.

Example ABR waveforms from pEEG and tEEG electrodes are shown in Fig. 2. Stimuli were FM-upsweeps at 0.5 ms duration, with harmonic structure and stimulus level as indicated. ABR waveforms are composed of a series of positive and negative peaks, with one or two sharp, prominent positive peaks in the range of 3-6 ms, followed by broader peaks extending up to 8-10 ms (Fig. 2). A prominent positive peak at a latency around 4 ms (asterisk) is visible in all recordings. Peak amplitudes are lower when recorded with tEEG than with pEEG electrodes, due to the nested analog differential processing used to achieve the common-mode noise cancellation function of tEEG. Waveform amplitudes were not normalized, as amplitude differences between electrode types or across recording days are not of interest in this initial study.

**Fig 2.**
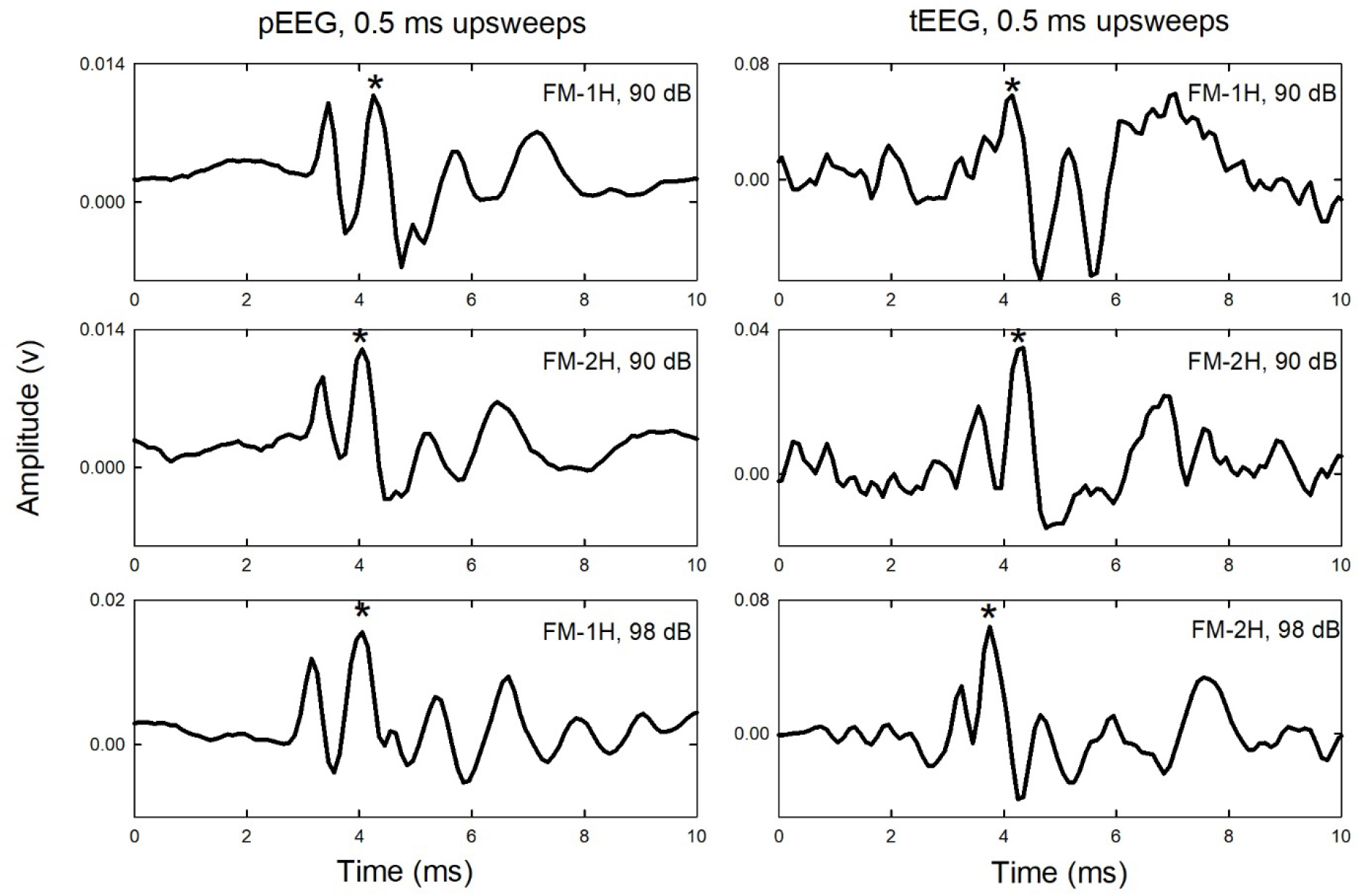
Example ABR waveforms recorded with pEEG electrodes (left column) and tEEG electrodes (right column). Stimuli are FM-upsweeps at 0.5 ms duration, with harmonic structure (FM-1H or FM-2H) and stimulus levels (90 dB peSPL, TDT tweeter; 98 dB peSPL, Kenwood tweeter) as indicated on the plots. pEEG data are from Bat J (top to bottom) 081519, 081519, and 062429. tEEG data are from (top to bottom) Bat J (012420), Bat J (012820), and Bat T (031419). The asterisk marks the second positive peak (latencies of around 4-5 ms) used for quantification of response amplitude for statistical testing (see Methods). Response amplitudes were not corrected for the different gains of pEEG and tEEG recordings.

ABRs to both FM-upsweeps and FM-downsweeps across sweep durations are shown in the form of heatmaps in Fig. 3 (data from three sample recordings). In these heatmaps, response amplitudes across the 10 ms post-stimulus interval are displayed on a color scale (brighter colors indicate stronger positive peaks, darker colors indicate stronger negative peaks). The heatmaps display the changes in amplitude of component peaks in the ABR with sweep duration and direction (x axes). Note that some ABRs (Fig 3A,B) show two initial prominent positive peaks; in these examples, both of these peaks are stronger (brighter colors) to FM-upsweeps compared to downsweeps.

**Fig 3.**
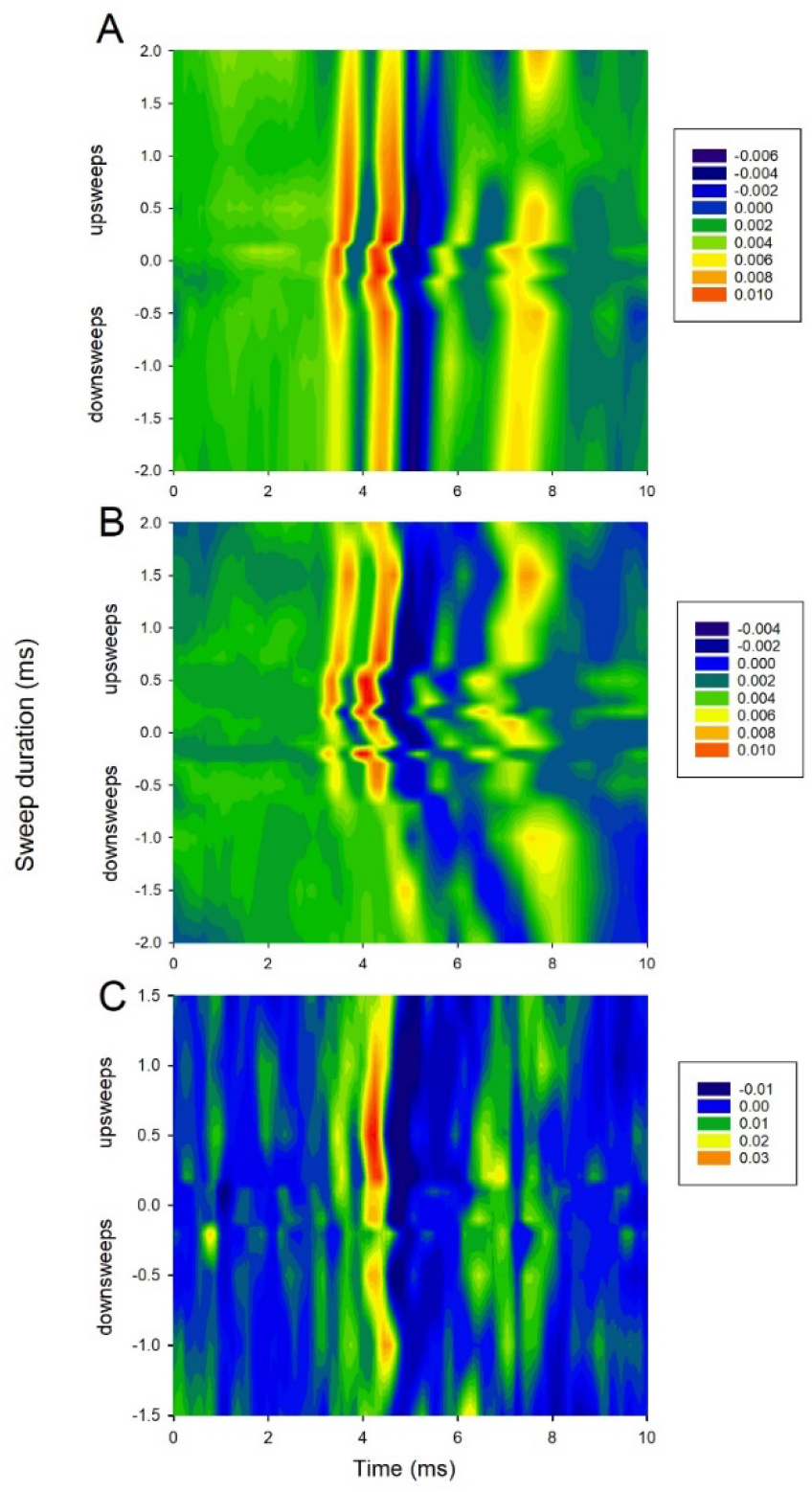
Heatmaps of responses to FM-upsweeps and downsweeps presented at 90 dB peSPL (TDT tweeter) across sweep durations (y axes). **A** pEEG responses from Bat J (081519), FM-1H sweeps. **B** pEEG responses from Bat J (081519), FM-2H sweeps. **C** tEEG responses from Bat J (012820), FM-2H sweeps. The upsweep duration producing the largest peak-to-trough amplitude (measured from the second positive peak; see Methods) is 0.5 ms up (**A**), 0.7 ms up (**B**), and 0.7 ms up (**C**).

To quantify the impact of FM sweep direction on the ABR, we measured the amplitude of the prominent positive peak within the latency range around 4 ms (asterisks in Fig. 2), to the subsequent negative peak (trough). This peak, usually the second positive peak in the ABR, was chosen because it was consistently visible in both pEEG and tEEG recordings, even though in some recordings (Fig. 3) the first positive peak was as high or higher in amplitude at some sweep durations. Peaks were measured only if they exceeded the calculated noise threshold values and only if the ABRs were not contaminated by bat movement. Amplitudes of the FM-up response are larger than those of FM-down response across the range of sweep durations tested (Fig. 4; data are from matched pairs in which both upsweeps and downsweeps were presented at the same sweep duration). Note that large variability (standard deviations) in response amplitudes. This variability stems from using absolute, rather than normalized, response amplitudes in the calculations. In addition, we found two comparisons in which the FM-downsweep elicited larger responses than the FM-upsweep at the shortest sweep durations of 0.1 and 0.2 ms.

**Fig 4.**
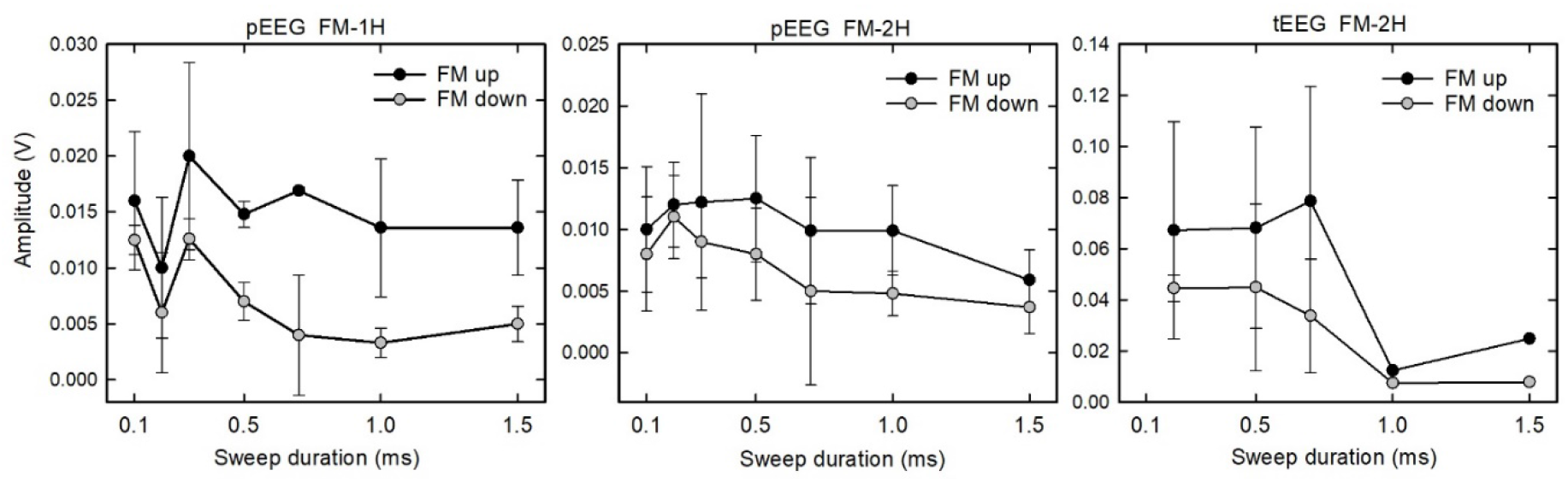
Peak-to-trough amplitudes vary with sweep direction and duration. Because only matched pairs of responses, where data are available for both upsweeps and downsweeps at the same sweep duration, are included, there is an unequal sample size across sweep durations. Data are plotted as means and standard deviations. Stimulus levels are 86, 90, or 98 dB peSPL, depending on the tweeter used. Not all recording days have the same number of matched pairs. Amplitudes are not normalized.

We compared peak-to-trough amplitudes to FM-upsweeps vs FM-downsweeps using two-tailed Wilcoxon signed-rank tests with Bonferroni correction of*p* values. We included in these analyses peak amplitudes at any stimulus level where matched pairs of responses to both upsweeps and downsweeps at the same sweep duration were available. Because the statistical tests were based on matched pairs, differences in stimulus parameters other than sweep direction (electrode type, sweep duration, harmonic content, stimulus level) do not affect the calculations. There was a statistically-significant higher peak-to-trough amplitude in response to FM-upsweeps compared to FM-downsweeps compiled across other experimental variables [*z* = 6.23, *df* = 82, *p* < 0.0001; Bonferroni critical value = 0.017]. We recalculated these tests for FM-1H and FM-2H sweeps separately. Results showed that FM-upsweeps evoked larger responses than FM-downsweeps for both stimulus types [FM-1H: *z* = 5.78, df = 46, *p* < 0.0001; FM-2H: *z* = 4.12, *df* = 43,*p* < 0.0005; Bonferroni critical value = 0.017]. We did not run statistical tests on data separated by electrode type, because the number of matched pairs for tEEG recordings was considerably smaller (*n* = 17) than those for pEEG recordings (*n* = 67). As shown in Figs. 3 and 4, however, the pattern of response is similar for both types of electrode recordings. We did not test statistically the effects of upsweep duration, because of unequal data points across durations.

We then asked whether an optimal duration of the FM-upsweep (i.e., that producing the largest peak-to-trough amplitude; Elberling et al. 2007) could be identified. As shown in Fig. 4, differences in response amplitudes across upsweep durations are small. The peak in the mean data for FM-1H sweeps is at 0.3 ms (pEEG electrodes, left plot). For responses to FM-2H sweeps, mean amplitudes in the duration range of 0.2 to 0.5 ms (pEEG electrodes, middle plot) are similar. The highest amplitude peak recorded by tEEG electrodes (right plot) occurred at 0.7 ms, but there are fewer data points represented in these analyses.

Finally, we examined whether responses to FM-upsweeps decreased in latency with decreases in stimulus level, as expected by amplitude-latency trading. Fig. 5 presents results from three pEEG recording sessions where stimulus level varied over at least a 20 dB level. Using the slopes for the 0 dB to 20-30 dB attenuations, the approximate amplitude-latency trading effect is 7-17 μs of added latency per dB attenuation. We were unable to collect sufficient data at a range of stimulus levels to quantify the presence of amplitude-latency trading in tEEG recordings.

**Fig 5.**
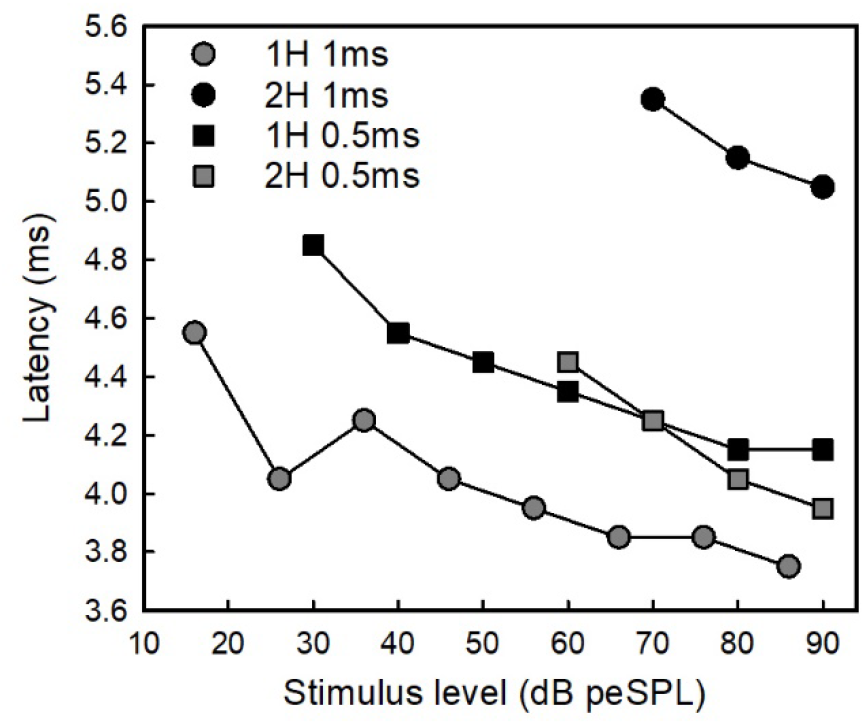
pEEG latencies to FM-upsweeps decrease with increases in stimulus levels, as expected from amplitude-latency trading. Data are plotted from four recording days in response to FM-1H and FM-2H upsweeps at two sweep durations, 1 and 0.5 ms. Data are from Bat J (legend order 062719, 091219, 091319, 091319).

## Discussion

Our goal in this experiment was to assess the feasibility of non-invasive scalp recordings of auditory-evoked activity in awake, unrestrained big brown bats. We tested two different electrode types, a conventional monopolar electrode and a novel tripolar electrode, on two female big brown bats who were trained to sit in a ceramic dish without excessive movements. For acoustic stimulation, we changed the duration and direction of FM sweeps within the frequency range of the bat’s echolocation sounds, in parallel with the stimuli used in our earlier recordings from the inferior colliculus (Luo et al. 2019). We hypothesized that peaks in the scalp-recorded ABR would follow the same pattern of response – higher amplitude responses to FM-upsweeps compared to FM-downsweeps – as observed in these earlier single and multiple unit recordings. In addition, we hypothesized that scalp-recorded ABRs would undergo amplitude-latency trading, as expected from invasive recordings. Our data support these hypotheses and thus verify the validity of our EEG recording techniques.

### ABR waveforms

ABRs are short latency responses reflecting synchronous activity along the auditory pathway from the eighth nerve up to the inferior colliculus (Picton et al. 1974). Results of invasive recordings from big brown bats indicate that peaks in the ABR have latencies that vary from 1-2 ms, reflecting activity from the eighth nerve, up to 4-6 ms, likely reflecting activity from the nucleus of the lateral lemniscus (Grinnell 1963; Suga 1969; Simmons et al. 1990; Haplea et al. 1994). From the cochlear nucleus to the inferior colliculus, however, single neurons exhibit progressively more latency variability, leading to response peaks with latencies extending up to 50 ms post-stimulus onset (Haplea et al. 1994; Ferragamo et al. 1998). This variability in response latency reflects the complexity of neural circuitry in the ascending auditory pathway and can make identification of the sources of the individual peaks in averaged ABR recordings challenging. The ABR peaks we quantified here have latencies within the range of 3.5-5.5 ms at the highest stimulus levels; these latency values are consistent with previous suggestions of an origin at the level of the nucleus of the lateral lemniscus (Suga 1969; Ferragamo et al. 1998; Boku et al. 2015). However, we cannot rule out contributions from other nuclei in the ascending pathway. Burkard and Moss (1994) recorded ABRs from needle electrodes inserted into the scalp of anesthesized big brown bats to FM-1H downsweeps of 1 ms duration. Response waveforms consisted of 4 peaks within the range of 2-6 ms at the highest stimulus level, with the most prominent positive peak (‘wave ib’ in their terminology) at around 2-3 ms. We observed in our data prominent peaks within this latency range, but this earlier peak was not always higher in amplitude than the subsequent positive peak. These differences in response latencies likely reflect the different stimulus envelopes used in the two studies. Burkard and Moss (1994) attributed a cochlear origin to an initial peak (‘wave ia’; mean latency of 1.16 ms) and indicated that later peaks originated at or below the level of the rostral pons and midbrain.

Both types of EEG electrodes used in our study also picked up broader peaks of activity at longer latencies up to 8-10 ms. Results of invasive recordings suggest that this longer latency activity originates from the inferior colliculus (Suga 1969; Simmons et al. 1990; Haplea et al. 1994). Because our focus was in comparing our data with those of Luo et al. (2019), we did not quantify these longer latency responses, but we note their presence. Our data show little evidence of longer latency (> 10 ms) responses that might reflect activity from the auditory cortex (Picton et al. 1974; Schalles et al. 2021). There are several reasons for this absence, including the placement of electrodes on the caudal half of the scalp and our choice of filter settings used to isolate the ABR. In addition, our experimental design was not reliant on the bat paying attention to or making choices between particular stimuli. These design factors are likely important in evoking longer latency ABR components that might reflect cognitive state (Picton et al. 1974; Schalles et al. 2021).

Scalp-recorded ABRs also showed evidence of amplitude-latency trading, at about 7-17 μs of added latency per dB attenuation. This trend of increasing latency with decreasing stimulus amplitude is expected from invasive recordings (Pollak 1988; Simmons et al. 1990; Klug et al. 2000) and from recordings with subcutaneous needle electrodes (Burkard and Moss 1994). This congruence in results serves to validate our recording technique. Simmons et al. (1990) reported an amplitude-latency trading ratio of 13–18 μs/dB from averaged local field potentials recorded from the inferior colliculus in response to FM-1H downsweeps; similar values (5-14 μs/dB) were found by Burkard and Moss (1994). Our values from scalp recordings are within these ranges.

### Relation to cochlear processing in bats

Our data are consistent with prior invasive recordings in big brown bats (Luo et al. 2019) and also with scalp recordings in bottlenose dolphins (Finneran et al. 2017) in showing that short latency ABRs are higher in amplitude to FM-upsweeps compared to FM-downsweeps of the same duration. The optimal duration of the FM-upsweep, that is, the duration producing the largest difference in response due to FM direction, has been proposed to provide an estimate of the speed of propagation of the traveling wave along the basilar membrane (Elberling et al. 2010). Luo et al. (2019) reported an optimal duration in the range of 0.5-1 ms in neural responses (local field potentials and multi-unit activity from three bats) recorded from the inferior colliculus to FM-1H sounds. Our data show that FM-upsweeps evoked larger scalp-recorded responses than FM-downsweeps, but with a mean optimal duration within the range of 0.3-0.7 ms, varying with stimulus harmonic content and electrode type. We note, however, that the differences in peak amplitudes recorded with scalp electrodes across the range of upsweep durations of 0.1-1.0 ms are small. For both optimal duration and comparisons between electrode types, we do not have enough data to determine if these differences reflect some biological function or simply reflect variability due to differences in sample size and quality of the scalp response. This is a topic for future research.

The concept of the optimal duration is based on modeling of basilar membrane mechanics of one-way traveling-wave propagation in a standard mammalian ear with no obvious species specializations (Dau et al. 2000; Elberling et al. 2007). In contrast, the ears of FM bats are specialized for ultrasonic hearing and echolocation. Specifically, in both big brown bats and Japanese house bats *(Pipistrellus abramus),* the stapes enters the cochlea, not at the base, but partway up the basal turn (Ketten et al. 2021). This unique anatomy suggests that simplified mammalian models of cochlear function based on one-way traveling wave direction of propagation may not apply to FM echolocating bats. As discussed recently (Shera 2022), these standard models may need to take into account standing wave reflections that depend on the location of active frequency tuning along the basilar membrane, which in bats could possibly be augmented by the placement of the stapes input further up the organ of Corti. These questions are best addressed through modeling efforts.

### Reliability and feasibility of noninvasive recordings in awake bats

The major goal of this study was to evaluate two methods for recording surface EEG signals from awake unrestrained bats. Both pEEG and tEEG electrodes recorded clear ABRs to FM sweeps. We had expected, given the design of the tripolar system for reducing movement artifacts, that higher quality signals would be picked up by tEEG. We observed that recordings from tEEG electrodes were of lower amplitude than those from pEEG electrodes and needed greater amplification; in addition, the placement of these electrodes on the scalp seemed more critical for evoking high quality responses. We were able to complete fewer tEEG than pEEG recording days due to various technical difficulties arising during the timeframe of these particular experiments. Because of these differences in sample size, the superiority of one technique over the other cannot be stated with certainty from our dataset. Subjectively, it appears that the ability of the unrestrained bat to remain motionless long enough to run through the entire experimental protocol was a stronger factor in collecting high quality data than the type of electrode. After training, the two bats did learn to sit relatively motionless in what to them seemed to be comfortable positions when electrodes were applied and stimuli presented, although in some cases we terminated experiments early if they began to make excessive movements. Further refinements to the technique might include using a different-sized or shaped dish in which bats sit during experiments, tailored to the individual bat’s preferences.

There are numerous important implications and interesting extensions that could follow from our demonstration of feasibility of noninvasive recordings in awake unrestrained bats. In some jurisdictions, including our own, collection of bats from the wild is strictly regulated due to conservation concerns and state wildlife capture limitations. Thus, non-invasive techniques such as we introduce here allow for reduction of numbers of animals needed for experiments as well as helping to maintain local wild populations. Importantly, noninvasive recordings from awake, unrestrained bats can be expanded to investigate perceptual and cognitive processing of biosonar signals, paralleling ongoing efforts in bottlenose dolphins (Schalles et al. 2021). Because in our experiments the animals are awake and unanesthetized, direct observation of attention and learning on echo identification, localization, and tracking is possible while recording simultaneously scalp-evoked activity, unlike in experiments involving anesthetized animals.

## DECLARATIONS

### Ethics approval

Brown University Institutional Animal Care and Use Committee

### Consent to participate

Not applicable

### Consent for publication

All authors agree to publication

### Availability of data and material

Data are available upon reasonable request to the first author, and will be uploaded to the Brown University data repository upon acceptance of the manuscript.

### Code availability

Code is available upon reasonable request to the first author.

### Authors’ contributions

AMS, AT, BMY, JAS designed research; AT, BMY, JAS performed research; AMS analyzed data; AMS, JAS provided equipment, space, and funding; AMS wrote the manuscript; all authors edited and approved the manuscript.

## Acknowledgements

A portion of these data was submitted by BMY as partial fulfillment of requirements for the Master’s degree in Biotechnology from Brown University. We thank Pedro Polanco for assistance with data analysis.

## Abbreviations

1H: one harmonic
2H: two harmonics
ABR: auditory brainstem response
EEG: electroencephalography
FM: frequency modulated
FM-downsweep: frequency modulated downsweep
FM-upsweep: frequency modulated upsweep
pEEG: EEG using pediatric electrodes
EEG: EEG using tripolar electrodes

